# Modelling seeding and neuroanatomic spread of pathology in amyotrophic lateral sclerosis

**DOI:** 10.1101/2021.02.24.431118

**Authors:** Sneha Pandya, Pedro D. Maia, Benjamin Freeze, Ricarda A L Menke, Kevin Talbot, Martin R Turner, Ashish Raj

**Affiliations:** Dept. of Radiology, Weill Cornell Medicine, 1300 York Avenue, New York, NY, USA; Wellcome Centre for Integrative Neuroimaging & Nuffield Department of Clinical Neurosciences, University of Oxford, Oxford, UK; Dept. of Radiology and Biomedical Imaging, University of California, San Francisco, USA

## Abstract

The neurodegenerative disorder amyotrophic lateral sclerosis (ALS) is characterized by the progressive loss of upper and lower motor neurons, with pathological involvement of cerebral motor and, additionally, extra-motor areas, in a clinicopathological spectrum with frontotemporal dementia (FTD). A key unresolved question is whether the distribution of pathology in ALS is driven by molecular factors such as regional gene expression, by differential network vulnerability, or is a combination of both. A system of histopathological staging of ALS based on the regional burden of TDP-43 pathology observed in *post mortem* brains has been supported to some extent by analysis of distribution of *in vivo* structural MRI changes. In this paper, computational modelling using a Network Diffusion Model (NDM) was used to investigate whether a process of focal pathological ‘seeding’ followed by structural network-based spread recapitulated *post mortem* histopathological staging and, secondly, whether this had any relationship to the pattern of expression of a panel of genes implicated in ALS across the healthy brain. Regionally parcellated T1-weighted MRI data from ALS patients (baseline n=79) was studied in relation to a healthy control structural connectome and a database of associated regional cerebral gene expression. The NDM provided strong support for a structural network-based basis for regional pathological spread in ALS, but no simple relationship to the spatial distribution of ALS-related genes in the healthy brain. Intriguingly, the critical seed regions for spread within the model were not within the primary motor cortex but basal ganglia, thalamus and insula, where NDM recapitulated aspects of the *post mortem* histopathological staging system. Within the ALS-FTD clinicopathological spectrum, non-primary motor structures may be among the earliest sites of cerebral pathology.

## Introduction

Amyotrophic lateral sclerosis (ALS), the commonest phenotype of motor neuron disease, is a progressive and fatal neurodegenerative disorder with complex molecular underpinnings (Talbot *et al*., 2018). The disease is clinically characterized by the progressive loss of upper motor neurons in the primary motor cortex and corticospinal tract and lower motor neurons of the spinal cord and brainstem. However ALS also involves extra-motor cerebral systems, with clear pathological, genetic and clinical overlap with frontotemporal dementia (FTD) (Es *et al*., 2017), with the behavioral variant being most common. Advances in neuroimaging have revealed many aspects of pathogenesis across the ALS-FTD spectrum (Turner *et al*., 2013; Chiò *et al*., 2014), with increasing interest in its potential to deliver therapeutic outcome measures (Menke *et al*., 2017).

The pattern of clinical symptom spread in ALS (Ravits *et al*., 2007b; Turner *et al*., 2010) and the associated spinal cord pathology (Ravits and La Spada, 2009), is not random but is focal in onset and spreads contiguously. Nearly all cases of ALS and around 50% of FTD are associated with cytoplasmic neuronal and glial inclusions of aggregated 43kDa transactive response DNA-binding protein, TDP-43 (Neumann *et al*., 2006). *Post mortem* histopathological classification has been interpreted as evidence of a stereotyped pattern of cerebral pathological involvement in ALS (Brettschneider *et al*., 2013). Several molecular mechanisms have been proposed to explain the apparent selective vulnerability of motor neurons. These include cell-autonomous factors, involving oxidative stress, excitotoxicity, and mitochondrial dysfunction (Turner *et al*., 2013) and non cell-autonomous factors involving cell-cell communication (e.g. glia (Philips and Rothstein, 2014)) or trans-neuronal transmission of aggregate-prone proteins through prion-like templating (Polymenidou and Cleveland, 2011; Riku, 2020), in which network connectivity might define the canonical pattern of spread (Seeley *et al*., 2009).

It is not yet clear how molecular vulnerability and network connectivity might combine in mediating regional patterns of pathology in ALS. Like other neurodegenerative diseases, the spatial topography of ALS histopathology is not related in a simple way to regional expression of genes implicated in pathogenesis (Fusco *et al*., 1999; Jackson, 2014; Subramaniam, 2019). A high level of clinical and molecular heterogeneity in the ALS-FTD syndrome meanwhile hamper the ability to map its course in a precise manner to facilitate effective therapeutic trials (Turner and Swash, 2015).

In this paper we address these issues using computational modeling, gene expression analysis and large observational imaging studies in ALS, combined with prior histopathological staging data. We interrogate whether focal seeding followed by structural network-based spread recapitulate patterns of ALS pathology by employing a Network Diffusion Model (NDM) to map neurodegenerative topography (Raj *et al*., 2012). This model was successful in recapitulating spatial patterns of diverse proteinopathies including Alzheimer’s (Raj *et al*., 2015), frontotemporal dementia (Raj *et al*., 2012), Parkinson’s disease (Freeze *et al*., 2018, 2019; Pandya *et al*., 2019), Huntington’s disease (Poudel *et al*., 2019) and progressive supranuclear palsy (Pandya *et al*., 2017).

## Methods

### Participants

Data used in this study were obtained after informed consent from participants in the longitudinal Oxford Study for Biomarkers in Motor Neuron Disease (‘BioMOx’) cohort based on referrals to a large tertiary ALS clinic and clinical assessment involving two experienced neurologists (KT, MRT). For the purposes of this group-level analysis, a diagnosis of ALS included those within all El Escorial clinical diagnostic categories at baseline (including those with pure upper or lower motor neuron syndromes clinically) who also showed clear progression of motor involvement in subsequent follow-up. Data were available for 79 such ALS participants (mean age at baseline 61±11 years, male:female 2:1, mean duration from symptom onset 49±57 months). Of these, 48 were also able to undergo repeat MRI every 6 months to a maximum of 5 visits in total (cohort overlaps with (Menke *et al*., 2018)). One baseline participant initially labelled as ALS was removed from the study cohort later due to failure to progress. All ALS participants were apparently sporadic (i.e. no family history of ALS or FTD). The study predated routine genetic testing in the clinic, but subsequent experience predicts this would have identified up to 3 apparently sporadic ALS gene mutation carriers, which is not felt to be material to the outcome.

### Image acquisition and regional volumetric analysis

Images were acquired using a 3T Siemens Trio scanner (Siemens AG) with a 12-channel head coil at the Oxford Centre for Clinical Magnetic Resonance (OCMR). A high resolution 3D MP-RAGE T1-weighted sequence was obtained for each subject with the following parameters: 192 axial slices; repetition and echo time (TR/TE)=2040/4.7 ms, flip angle 8°, 1 × 1 × 1 mm^3^ voxel size, and 6 min acquisition time.

68 cortical and 18 subcortical volumes from these MRI images were extracted using FreeSurfer software with a cross-sectional pipeline for both the cohorts. Regional volumes were normalized by total intracranial volume generated by FreeSurfer to correct for head size. Image processing steps were visually inspected for white–gray matter boundary and skull-stripping errors to ensure they had been carried out correctly. 6 subjects that rated either ‘partial’ or ‘fail’ due to FreeSurfer failure or insufficient tissue contrast were excluded from analysis. A vector of regional atrophy was created by using a two tailed *t*-test between ALS and normal mean ICV corrected regional volumes such that *t_ALS_ = {t_ALS_(i)|i ∈ [1,N]} (N = 86)*. The *t*-statistic was converted to the natural positive range between 0 and 1 using the logistic transform given by Φ = 1/(1+exp(-(t_ALS_ – a0)/ *σ* /std(t_ALS_)), where, *σ* = 2 and a0 = 0.5* *σ* (Raj *et al*., 2015). This transformation maps t-values such that they asymptotically approach 0 as *t*_ALS_ approaches −∞ and 1 as *t*_ALS_ approaches +∞. The parameter *σ* controls the steepness of the logistic function. These atrophy measures were then used to test the propagation modeling analyses using NDM.

### A healthy structural connectome

Axial T1-weighted structural MRI scans using fast spoiled gradient-echo sequence(TE = 1.5 ms, TR = 6.3 ms, TI = 400 ms, 15° flip angle, 230 × 230 × 156 isotropic 1 mm voxels) and high angular resolution diffusion tensor imaging data (DTI) (55 directions, b = 1,000 s/mm2, 72 1.8-mm-thick interleaved slices, 0.8594 mm × 0.8594 mm planar resolution) were acquired on a 3T GE Signa EXCITE scanner from fully consented 73 young healthy volunteers under a previous study approved by Weill Cornell’s institutional review board (Kuceyeski *et al*., 2013). Thus, the cohort used to extract a healthy structural connectome was different from the 38 age-matched controls described earlier, that was used for determining regional atrophy. Probabilistic tractography was performed on the diffusion MRI data after seeding each voxel at the interface of the WM and GM boundary. The resulting streamlines were binned into subsets corresponding to every pair of GM regions given by the parcellation scheme described above. The anatomical connection strength (ACS), a measure of connectivity, was used in this paper (Iturria-Medina *et al*., 2007). ACS is defined as the weighted sum of the streamlines found to exist between any pair of gray matter structures, weighted by each streamline’s probability score. The ACS is further normalized by a scaling factor equaling to the total sum of all streamlines. We define *c_i,j_* as the resulting connection strength between *i*^th^ and *j*^th^ GM regions. We refer to the matrix collecting all pair-wise entries as the connectivity matrix *C = {c_i,j_}*. Here the ACS is used as an approximation of the cross-sectional area of all axonal projections connecting two regions – a plausible choice given our goal of modeling the amount of pathology transmission conducted through these projections. Connections are assumed to be bidirectional since directionality is not deducible from DTI tractography data.

### NDM for ALS pathology spread

The hypothetical spread of disease-causing proteinopathy into the network represented by the connectivity matrix *C* over time *t* can be captured by starting a diffusion process from a “seed” region. Since we do not know *a priori* which region is the likely seed, we select every brain region as the seed region, one at a time (Raj *et al*., 2012). The overall strategy is to simulate a diffusion process on the connectivity graph for many time points, starting from each seed location, while recording its correlation with measured regional atrophy maps.

We modeled ALS progression as a diffusion process of the pathology load x on the graph *C* over model time *t*. From (Raj *et al*., 2012) the transmission of pathology from region 1 to region 2 is represented as 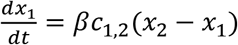 where *x_1_* and *x_2_* denote the extent of disease-causing pathology in each region, and *β* is a global diffusivity constant. Denoting pathology from all regions *i* into a vector **x**(t) = *{x_i_(t)}*, the above equation extends to become:

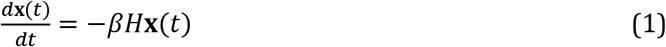

*where H* is the well-known graph Laplacian

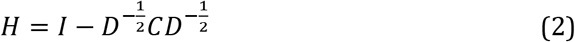

where D is a diagonal matrix whose diagonal entries contain the degree of each node, degree being defined as the sum of weighted connections emanating from the node. Note, in order to accommodate regions having widely different out-degrees, we used the degree-normalized version of the Laplacian matrix (Raj *et al*., 2015). Eq. 1 admits a closed-form solution **x**(t) = *e^−βHt^***X_0_** where **x_0_** is the initial pattern of the disease process at *t* = 0, and we call term *e^—βHt^* the *diffusion kernel* since it acts essentially as a spatial and temporal blurring operator on **x_0_**. The unit of the model’s diffusion time *t* is arbitrary (au). Global diffusivity *β* is unknown, hence we chose a value that would roughly span ALS progression (3-10 years), giving *β* = 1.

The NDM is described by pathology x(t) and our hypothesis is that it should correlate with empirical atrophy Φ, Pearson correlation strength (R statistic) and p-values were calculated between the (static) empirical atrophy measured on the ALS group Φ and x(t) at all model timepoints t.

### Repeated seeding

The NDM was run for all 86 seed regions, each time starting from a different ROI, such that **x_0_** is a unit vector with 1 at the index of the seed and zeros at all other regions. We observed that the atrophy pattern in our group was generally bilateral, hence for repeated seeding experiments, we chose to seed bilaterally, so that two entries in the “unit” vector were assigned 1. This was repeated for each region in turn, and the NDM-predicted pathology pattern was calculated. For each predicted pathology vector *x^i^(t)* seeded at region i, the Pearson’s correlation coefficient R occurring over all model timepoints *t* was determined, giving *R^i^(t)*. “R-t curves” were represented by plotting these *R^i^(t)* values on common axis. 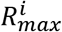 was recorded as the maximum value from each *R^i^(t)*, which reflected the likelihood of i-th region as the true region of pathology onset.

### Histopathological staging

We tested whether NDM recapitulated the *post mortem* histopathological staging published in ALS (Brettschneider *et al*., 2013), assigning each of the 86 regions available in our atlas an ALS stage from 1 to 4. Regions that were not part of this staging schema were arbitrarily assigned stage 5, a category that denotes the least vulnerable regions X.

### Spatiotemporal evolution from most likely seed regions

The top five regions *i* with the highest 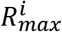 were chosen to qualify as best seeds, along with the precentral gyrus which is considered a key early site of ALS. We tabulated a similar list of top 5 seed regions that gave the highest *R_max_* (Pearson correlation) between the NDM and the ALS TDP-43 pathology staging data. An average of these two *R_max_* values was computed to give a representation of R (*R_avg_*) from both empirical atrophy and TDP-43 staging. The regions with the highest *R_avg_* were considered as the best seeds overall and were used as seeding locations for further analysis (see Results).

### Regional gene expression analysis

Prominent genes linked to familial ALS were (n=25) identified from various studies (Robberecht and Eykens, 2015; Smith *et al*., 2017; Vajda *et al*., 2017; Chia *et al*., 2018; Karch *et al*., 2018; Nicolas *et al*., 2018) and mapped to 86 regions in the Desikan-Killiany atlas as in (Freeze *et al*., 2018). Additionally, genes in which pathogenic variants have been associated with TDP-43 pathology (n=26) (Scotter *et al*., 2015) were also mapped to 86 regions as above. A list of all the genes used in this study can be found in the supplementary data. For each gene, data was obtained from the publicly available human Allen Brain Atlas (ABA) (Hawrylycz *et al*., 2012). Briefly, the ABA includes 926 brain regions, with each region having microarray expression levels from a set of 58,692 probes that correspond to 21,245 distinct genes. Expression data for each of the 926 regions from ABA were mapped to the 86 regions of the Desikan-Killiany atlas. All samples for all probes within the same region were averaged and then normalized for each gene to produce a single expression value quantified as a z-score. White matter tracts were excluded from the analysis. Expression for each gene was averaged for six subject brains (which comprises data for 6 left hemispheres and 2 right hemispheres; more information can be found at help.brain-map.org/download/attachments/2818165/Normalization_WhitePaper.pdf).

### Statistical analyses

Throughout this paper the primary test statistic used to evaluate all models was Pearson’s correlation strength R. In each case the dependent variable was the vector of regional atrophy or ALS staging, while the dependent variables were the NDM-predicted regional vector, and/or regional gene expression. As described above, R was computed at each model time t and the highest value was chosen as the model evidence.

For the gene results, genes for each category were corrected for multiple comparisons using Bonferroni method, with thresholded p_corr_ = 0.05/25 = 0.002 for ALS-related genes and thresholded p_corr_ = 0.05/26 = 0.0019 for TDP-43 specific genes. Correlation coefficients with p-values less than p_corr_ were considered statistically significant. Next, L_1_ regularized regression model was created containing NDM from the seed region at *t_max_* 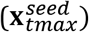 and regional genetic expression profiles averaged across all subjects and probes. Ten-fold cross-validation was performed for each model across a range of values for the tuning parameter lambda (λ) using the Matlab script ‘lasso’. These mapped genes and 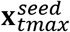 were then correlated with the atrophy to achieve significant predictors for atrophy. Predictors for each lasso were corrected for multiple comparisons using Bonferroni method, with thresholded 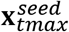 ALS-related genes, and TDP-43 specific genes. Correlation coefficients with p-values less than p_corr_ were considered statistically significant.

### Random scrambling

In order to build a null distribution for assigning significance to the NDM, we performed two levels of randomization experiments. 1) We ran the NDM on 2000 randomly scrambled versions of the connectivity matrix C. C was scrambled using a symmetric transformation of the network’s nodes by randomly permuting entire rows and columns, and the NDM was evaluated for each shuffled network after bilateral Insula seeding. This scrambling procedure maintains the edge and node statistics of the true connectivity C. The NDM evaluated on these 2000 randomly scrambled networks therefore constitute null or reference models which supplied significance values to results of the true model. 2) We ran the NDM on 2000 randomly scrambled ALS atrophy vector. Atrophy values in *t_norm_* vector were randomly assigned amongst the 86 cerebral regions with 2000 different permutations. This scrambling method maintained the true connectivity C but replaced true regional atrophy pattern with a random distribution of atrophy.

### Data availability

All data used in this study will be made available upon reasonable request and relevant code will be uploaded to https://github.com/Raj-Lab-UCSF repository. Oxford’s Wellcome Centre for Integrative Neuroimaging has an inherent commitment to data-sharing. To get access to the data and comply with the research ethics committee approval an application to the corresponding author will be required so that the precise geographical extent of sharing is known.

## Results

### Spatial distribution of ALS atrophy and repeated seeding of the NDM

Figure 1A shows glass-brain (LoCastro *et al*., 2014; Marinescu *et al*., 2019) illustrations of spatial distribution of ALS atrophy from our cross-sectional cohort which is consistent with progression of ALS pathology (Kassubek *et al*., 2005; Grosskreutz *et al*., 2006; Mezzapesa *et al*., 2007; Agosta *et al*., 2010; Westeneng *et al*., 2015). Pathology in each region is proportional to the t-statistic of ALS atrophy after logistic transform, where color towards red show increased severity. Table SI-1 shows empirical atrophy values of top 20 regions averaged across both the hemispheres.

**Figure 1:**
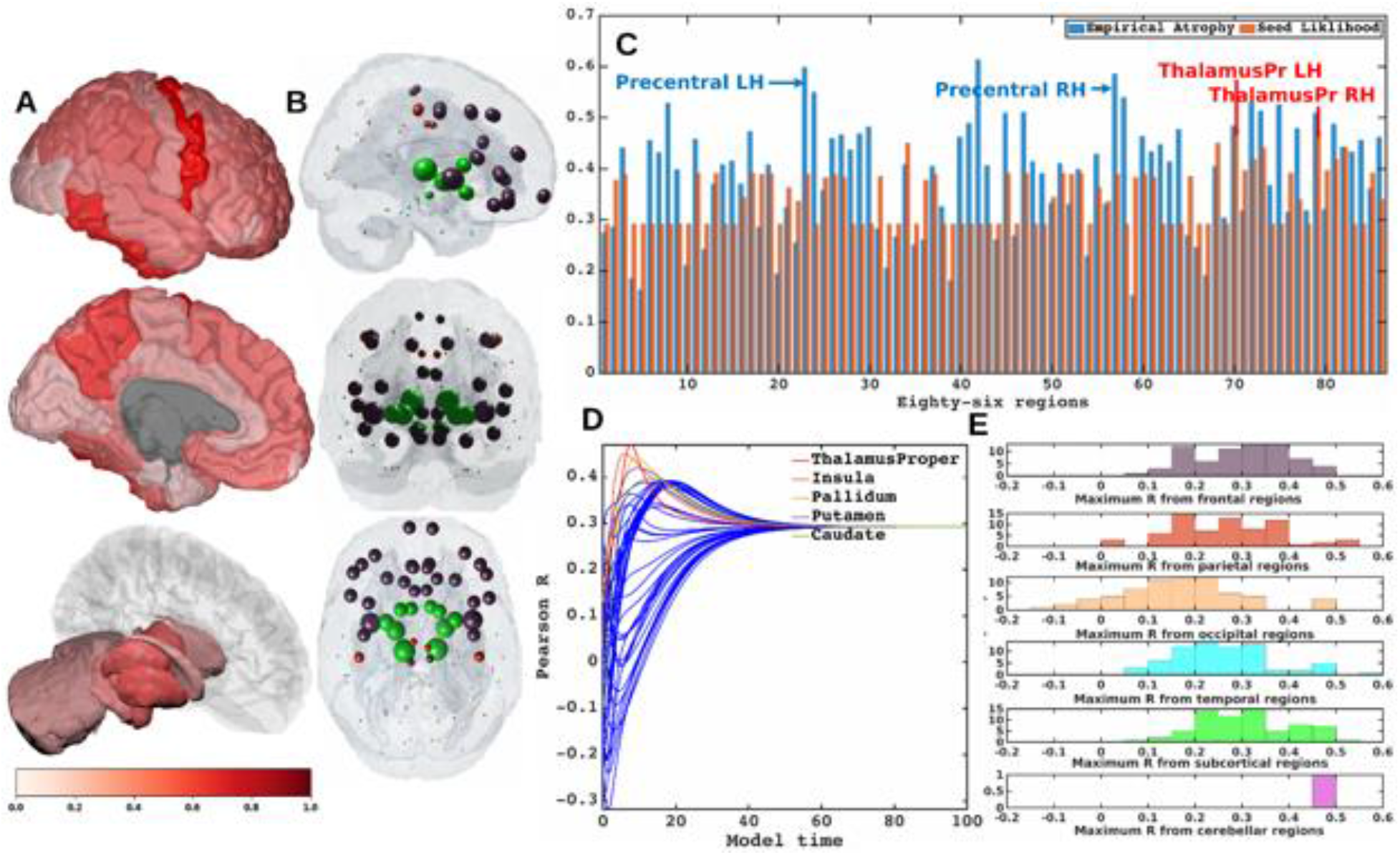
Spatial distribution of ALS atrophy and repeated seeding. A] Measured regional ALS atrophy are depicted by glass brain visualization. Bilateral volume loss was observed in somatosensory, frontotemporal, and subcortical regions, with most atrophy occurring in precentral gyrus, inferior temporal gyrus, precuneus, putamen, and thalamus regions. Severity of disease in each region is depicted in a color bar, where color towards red show increased severity. B] Each region was seeded in turn and NDM was played out for all time points. Pearson’s R was recorded at each time point between the model and ALS atrophy vector. As the diffusion time increases, more and more of the pathogenic agent escapes the seed region and enters the rest of the network. The point of maximum correlation with measured atrophy was recorded with glass brains of measured R with spheres placed at the centroid of each brain region, and their diameter proportional to effect size. Spheres are color coded by lobe – frontal = purple, parietal = red, occipital = orange, temporal = cyan, subcortical = green, and cerebellum = magenta. C] Histogram of empirical atrophy and seed region likelihood as represented by R_max_ is shown side-by-side. Precentral which is the highest atrophied region when taking the average of empirical atrophy from left hemisphere (LH) and right hemisphere (RH) is not the best seed, thereby suggestive of inconsequential role of higher atrophy values in determination of R_max_. D] NDM seeded at bilateral regions indicates that the thalamus is the one of the most plausible candidate for ALS seeding – it has the highest peak R, and the characteristic intermediate peak indicative of true pathology spread. Other regions among the top five that obtained the highest R were insula, pallidum, putamen, and caudate. R-t curves for the remaining regions are shown in blue. E] Histogram of maximum R achieved from six major regions for all individual subjects. Rmax values were attained for each of these regions from 79 individual subjects. We can see that for most of the subjects’ maximum R (Rmax > 0.4) was achieved from the frontal and subcortical regions compared to other regions.

Each region was computationally seeded in succession and the NDM evolved over model time *t* on the healthy connectome *C*. The spatial distribution of *R_max_*, which is indicative of the likelihood of each region as a seed is depicted (Figure 1B). Figure 1C shows the distribution of empirical atrophy and *R_max_* for each of these 86 regions alongside, which indicates that the NDM-derived seeding propensity value (R_max_) does not simply reproduce the regions displaying the highest atrophy but instead reflects the consequence of network transmission starting from that region.

Figure 1D shows the R-t curve revealing spread of *R_max_* corresponding to the best fit between empirical data and the NDM seeded at the i^th^ region (Eq. 2). For each region, the R-t curve would yield an intermediate peak in R, resembling the best match between the NDM and empirical data, and then after diffusing uniformly with decreasing resemblance between the actual data and the NDM. Table SI-2 shows top 20 regions with maximum Pearson correlation strength for each region seeded in succession. Basal ganglia structures and the insula were among the top best seed regions for predicting ALS atrophy (Table SI-2), with the insula serving as the most likely seed region for ALS with the highest *R_avg_* as seen in Table 1.

**Table 1:** List of regions most likely to serve as seeds of ALS pathology. Seeding likelihood of a region, shown in the rightmost column, is denoted by the average of the highest R from both empirical atrophy and histopathological staging. Although the precentral gyrus does not have the highest seeding likelihood by these measures, it was included here due to its prominence as an early site in ALS.

| Region | Atrophy | $R_{\max}$ (NDM vs Atrophy) | R (NDM vs ALS Staging) | Average R |
| --- | --- | --- | --- | --- |
| Insula | 0.41 | 0.45 | -0.64 | 0.55 |
| Putamen | 0.51 | 0.42 | -0.65 | 0.53 |
| Palladum | 0.48 | 0.44 | -0.61 | 0.53 |
| Caudate | 0.32 | 0.40 | -0.52 | 0.46 |
| Precentral | 0.59 | 0.39 | -0.42 | 0.40 |
| Thalamus | 0.50 | 0.47 | -0.30 | 0.38 |
| $R_{\max}$ – Maximum Pearson Correlation | | | | |

To test whether the above results on group regional data and generic ALS staging data are also applicable to individual ALS subjects, we also ran NDM on ALS individuals and calculated maximum Pearson’s R after seeding each of 43 bilateral ROIs for each subject. Peak R is achieved by frontal, parietal, temporal and subcortical regions, with frontal and subcortical regions achieving *R_max_* > 0.4 from most subjects (Figure 1E). Thus, there is considerable inter-subject heterogeneity in seeding, while at the group level there is a convergence of the likely seeds in frontoinsular and BG regions.

### Comparison with histopathological staging

Table 2 shows ALS staging for TDP-43 pathology in each region. As shown in Figure 2 MRI atrophy and ALS staging are not highly correlated (R = −0.27), hence we wished to assess whether our model is also able to recapitulate *post mortem* histopathological staging. Therefore, the most likely seeding location for NDM was determined based on criteria that worked best for both atrophy and histopathological staging. Based on this criterion, the insula was selected as the best seed and used to play out NDM for all subsequent analyses (Table 1). Empirical atrophy, 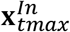 and staging maps are shown in Figure 2A, 2B, and 2C respectively. Figure 2B shows the distribution of predicted atrophy as determined by 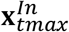. Pathology severity in each region is proportional to the color gradient. Regions with maximum severity as indicated by the NDM corresponded to regions in the more advanced histopathological stages. Given that seeding from the thalamus (Th) consistently produced the best R against empirical data (Figure 1D, Table SI-2), results were also obtained from 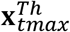 (Figure SI-1). This demonstrated bilateral volume loss mainly occurring in regions corresponding to advanced histopathological staging.

**Figure 2:**
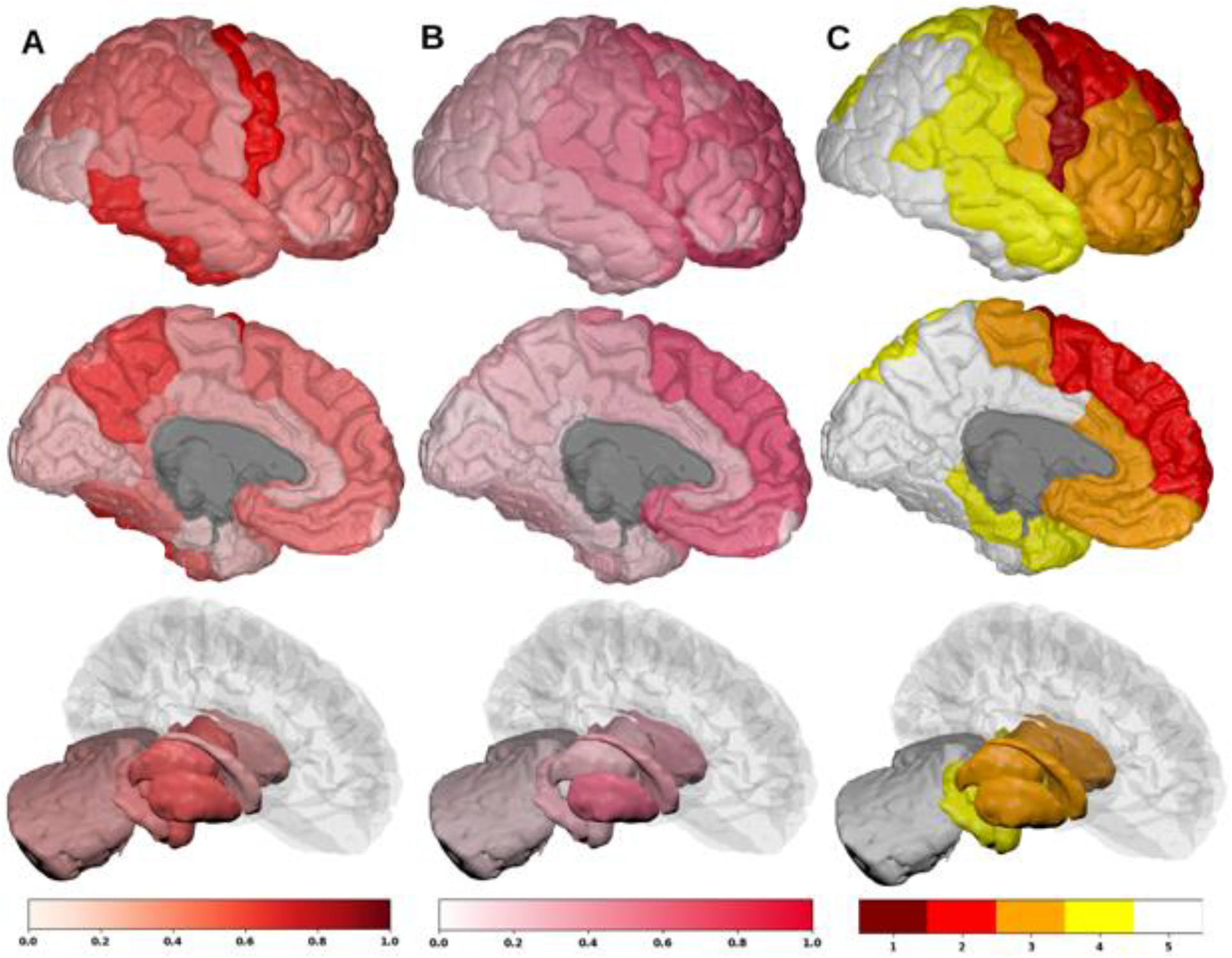
Spatial distribution of ALS atrophy, NDM Predicted atrophy and histopathological staging. A] Measured regional ALS atrophy are depicted by glass brain visualization. Bilateral volume loss was observed in somatosensory, frontotemporal, and subcortical regions, with most atrophy occurring in precentral gyrus, inferior temporal gyrus, precuneus, putamen, and thalamic regions. Severity of disease in each region is depicted in a color bar, where color towards red show increased severity. B] Glass brains of NDM seeded at the bilateral insula at t_max_ = 6.06 au yields progression of ALS from insula to connected, subcortical, anteromedial portions of temporal lobe and frontal areas. Bilateral volume loss is mainly observed in frontal and subcortical regions, with most atrophy occurring in later orbito-frontal, superior frontal, precentral, rostral middle-frontal, and putamen regions. Severity of disease in each region is depicted in a color bar, where color towards magenta showing increased severity. C] The ALS stage from 1-4 for each of the 43 bilateral regions. Stage 1 (maroon) starts with agranular motor cortex. The next affected regions (stage 2 in red) are the premotor cortex and parts of prefrontal neocortex. The pathology then progresses into striatum and into the prefrontal/postcentral cortices (stage 3 in orange), and finally to stage 4 show (yellow) involving anteromedial portions of the temporal lobe and the hippocampus. Stage 5 shows regions in white that are not part of the published histopathological staging system.

**Table 2:** Post mortem histopathological stages for each of 43 bilateral regions

| Region Names | ALS Staging | Region Names | ALS Staging | Region Names | ALS Staging |
| --- | --- | --- | --- | --- | --- |
| Bankssts | 5 | Parsorbitalis | 3 | CerebellumCortex | 5 |
| Caudalanteriorcingulate | 3 | Parstriangularis | 3 | Thalamus_Proper | 3 |
| Caudalmiddlefrontal | 2 | Pericalcarine | 5 | Caudate | 3 |
| Cuneus | 5 | Postcentral | 3 | Putamen | 3 |
| Entorhinal | 4 | Posteriorcingulate | 5 | Pallidum | 3 |
| Fusiform | 5 | Precentral | 1 | Hippocampus | 4 |
| Inferiorparietal | 5 | Precuneus | 5 | Amygdala | 4 |
| Inferiortemporal | 5 | Rostralanteriorcingulate | 3 | Accumbensarea | 3 |
| Isthmuscingulate | 5 | Rostralmiddlefrontal | 3 | Hypothalamus | 4 |
| Lateraloccipital | 5 | Superiorfrontal | 2 |  |  |
| Lateralorbitofrontal | 3 | Superiorparietal | 4 |  |  |
| Lingual | 5 | Superiortemporal | 4 |  |  |
| Medialorbitofrontal | 3 | Supramarginal | 4 |  |  |
| Middletemporal | 4 | Frontalpole | 3 |  |  |
| Parahippocampal | 4 | Temporalpole | 4 |  |  |
| Paracentral | 3 | Transversetemporal | 4 |  |  |
| Parsopercularis | 3 | Insula | 3 |  |  |

### Spatiotemporal evolution of ALS atrophy

The spatiotemporal evolution of ALS atrophy as recapitulated by network diffusion and evolved from the insula at model times *t* = 2,4,6 (*au*) is shown in Figure 3. The evolution of network diffusion process seeded at the insula starts at early stage (t=2) through mature stage (t=6), where the maximum correspondence of NDM to empirical data occurred. Here, time is arbitrary, hence we have used *“au”* as the unit of time for illustrative purpose only. At the initial stage, the disease involves subcortical and frontal regions, followed by motor regions, and finally showing widespread involvement of extra-motor and cortical regions. A linear positive association between empirical atrophy and predicted atrophy (R = 0.45, p_corr_ < 5.8×10^−4^) as represented by the NDM at model times *t* = 2,4,6 (*au*) was demonstrated (Figure 3B). A linear association was found between the histopathological staging and the NDM at model times *t* = 2,4,6 (*au*) (Figure 3C), with a negative correlation between the predicted atrophy and stage, which as model time progresses, shows increased correlation between the NDM prediction from insula-seeding and each stage (R = −0.66, p_corr_ < 5.8×10^−4^). Similarly, we also explored spatiotemporal evolution from thalamic-seeding at model times *t* = 3,5,8 (*au*) (Figure SI-2). With thalamic-seeding, the predicted disease course involved mainly subcortical regions at the initial stage, followed by diffusion into the frontal and motor regions.

**Figure 3:**
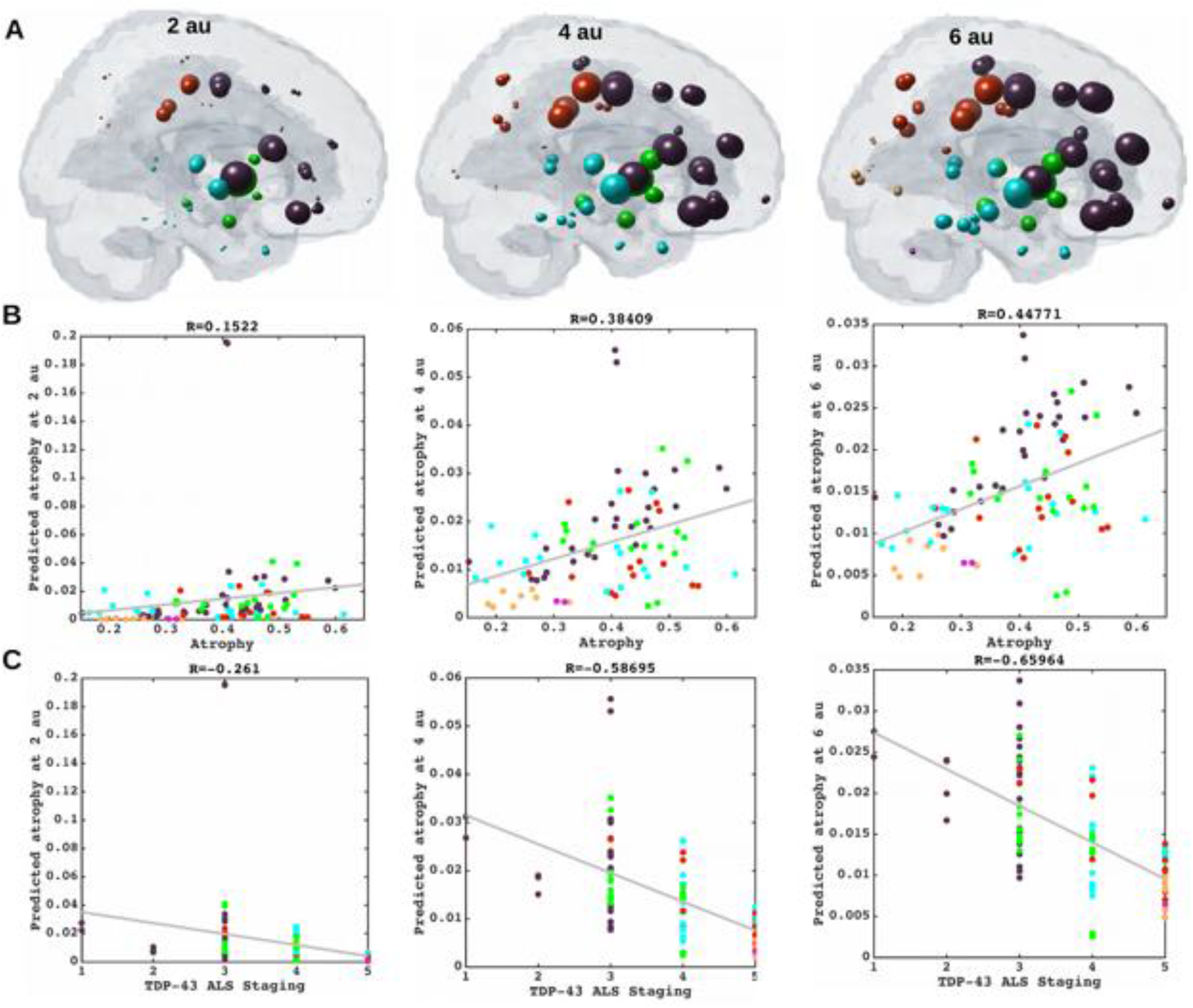
Spatiotemporal evolution, scatter plots of atrophy and histopathological staging with NDM at different model times. A] Evolution of insula-seeded network diffusion at model times *t* = 2,4,6 (*au*) exhibited frontal involvement initially, followed by slower diffusion into the temporal, motor and subcortical cortices, and finally showing widespread involvement of cortical regions. This temporal sequencing predicted by the model suggest that volume loss in ALS involves extra-motor regions, particularly the prefrontal and subcortical regions. B] Scatter plot of NDM from insula versus empirical ALS atrophy at model times *t* = 2,4,6 (*au*). Dots are color coded by lobe - frontal = purple; parietal = red; occipital = orange; temporal = cyan; subcortical = green; and cerebellum = magenta. A positive correlation is observed between ALS empirical atrophy and NDM predicted atrophy from bilateral insula, which increases significantly (R = 0.45, p_corr_ < 5.8×10^−4^, 0.05/86) at matured model times *t* = 4,6 (*au*). C] Scatter plot of NDM from insula versus histopathological staging at model times *t* = 2,4,6 (*au*). Dots are color coded by lobe - frontal = purple; parietal = red; occipital = orange; temporal = cyan; subcortical = green; and cerebellum = magenta. A negative correlation was observed between the NDM and ALS staging from bilateral insula, which decreases significantly (R = −0.66, p_corr_ < 5.8×10^−4^) at matured model times *t* = 4,6 (*au*). As time progressed, greater frontal, temporal and subcortical regions were involved with NDM closely resembling empirical ALS-FTD pathology.

### Relationship of atrophy to regional ALS risk gene expression

The linear relationship between different categories of genes (listed in Table SI-3) and empirical atrophy was studied (Figure 4). Figure 4A shows distribution of empirical atrophy for reference. Figures 4B and 4C show scatter plots of empirical atrophy versus averages of ALS-related genes, and TDP-43 specific genes respectively. No association was found between ALS-related genes, TDP-43 specific genes empirical atrophy. Further, no association was found between atrophy and TARTDP gene itself (4D), which codes for TDP-43. Figures 4E, 4F, and 4G show local distribution of average of ALS-related genes, average of TDP-43 specific genes and TARDBP, respectively. Table SI-3 shows correlations of ALS-related genes, and their PCA vs. empirical atrophy (to the left) and correlations of TDP-43 specific genes and their PCA vs. empirical atrophy (to the right). These results suggest that genes alone do not contribute to regional vulnerability, and that linear association between atrophy and gene expression profiles are complex and cannot be explained by univariate analysis of genes alone.

**Figure 4:**
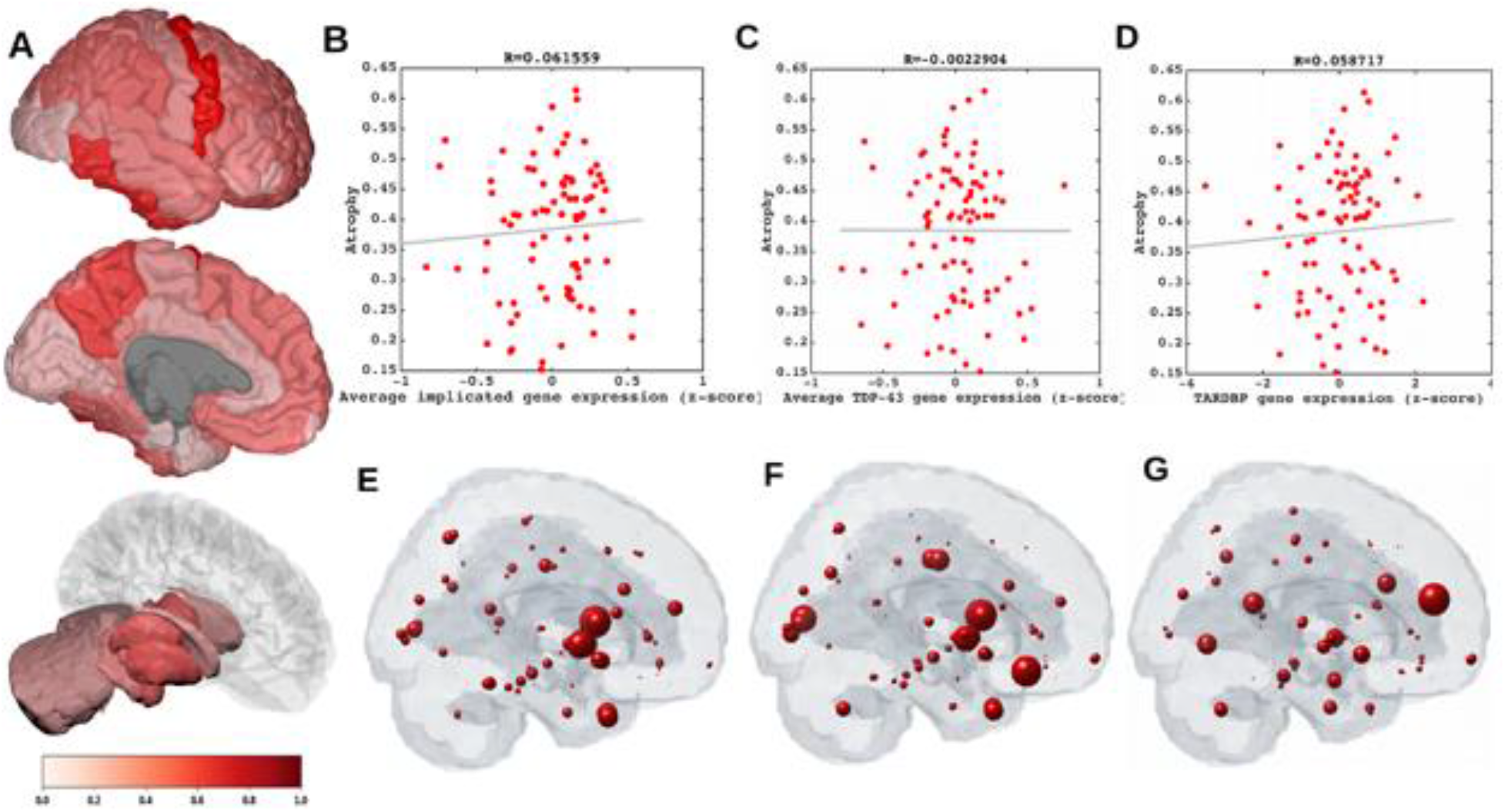
Spatial distribution of ALS atrophy, scatter plots of genes vs ALS atrophy, spatial distribution of genes. A] Measured regional ALS atrophy are depicted by glass brain visualization. Bilateral volume loss was observed in somatosensory, frontotemporal, and subcortical regions, with most atrophy occurring in precentral gyrus, inferior temporal gyrus, precuneus, putamen, and thalamic regions. Severity of disease in each region is depicted in a color bar, where color towards red show increased severity. B] Scatter plot of empirical atrophy vs average of all ALS-related genes shows no clear association. C] Scatter plot of empirical atrophy vs average of TDP-43 associated genes (F) shows no clear association. D] Scatter plot of empirical atrophy vs TARDBP gene expression (G) also shows no clear association – this was chosen for comparison, given that a small minority of ALS cases involve mutations in TARDBP. Spheres in glass brains were placed at the centroid of each brain region, and their diameter was proportional to effect size.

### NDM atrophy and regional ALS risk gene expression

Although genes do not bear an association with regional atrophy directly, it is possible that they may contribute to regional atrophy along with network transmission of pathology. To test this, we used cross-validated L_1_ regularized regression (LASSO) feature selection to identify highly significant predictors of ALS pathology from the NDM and ALS-related genes, and those genes that might result in TDP-43 misfolding via downstream events. The model included the NDM predictor 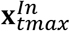, 25 ALS-related genes, and 26 TDP-43 specific genes (Figure 5). Six predictors survived lowest MSE in our model as seen in Figure 5: 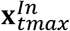**, CCNF*, UBQLN2*, PFN1*, and CSNK1E* with insula-seeding, suggesting that the NDM is the best predictor of ALS atrophy, and that only few genes in addition to the NDM contribute to the spatial pattern of disease, but in a weaker capacity.

**Figure 5:**
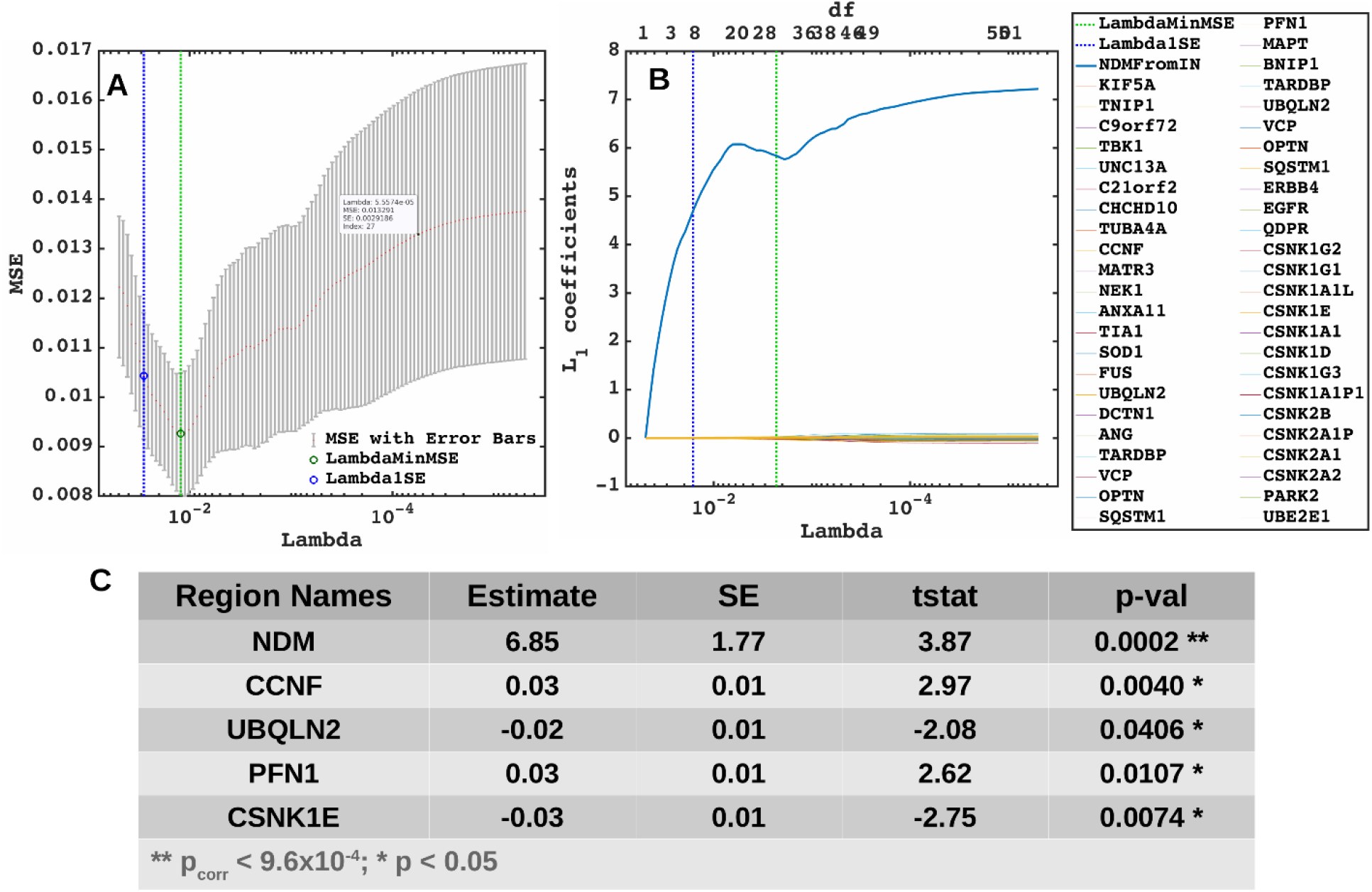
Lasso plots and model parameters. A] Ten-fold cross validated MSE curves for determining regularized parameter lambda. Predictors with minimum L_1_ coefficient as a function of regularized parameter lambda with no more than one standard deviation (blue dotted line) were considered to be the most favorable. B] Cross-validated L_1_ regularized regression coefficients as a function of tuning parameter lambda for a model containing the NDM from insula, ALS-related genes, and genes implicated in trans-synaptic TDP-43 transfer as predictors. Trace plot shows that as lambda increases towards the left, lasso sets various coefficients to zero, thereby removing them from the model. C] Model parameters and p-values of significant predictors that survived with p < 0.05 (represented with “*”) and with Bonferroni corrected p (represented with “**”). The NDM and expression profiles of CCNF, UBQLN2, PFN1, and CSNK1E have non-zero coefficients at minimum model MSE, indicating that these are essential predictive variables.

Given that thalamic-seeding achieved the highest R from empirical data, LASSO analysis was repeated with NDM from thalamic-seeding using 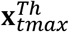 and different categories of genes. With thalamic-seeding of NDM, 10 predictors survived: 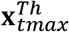**, KIF5A*, TBK1*, TIA1*, BNIP1*, CSNKIG1*, CSNK2A1*, CSNK2A1P*, and CSNK2A2* (Figure SI-3). However, again genes were far less significant contributors than NDM.

### NDM evaluation against alternate modelling of network connectivity and ALS atrophy

To test the predictive power of NDM against alternate network models, we evaluated its specificity to ALS atrophy and to the connectome upon which it evolves. The distribution of Pearson’s R over 2000 randomly simulated connectome matrices and atrophy vectors from insula and thalamic-seeding are shown in Figure 6 and Figure SI-4 respectively. Random model’s R was much lower than the maximum R of 0.45 from insula-seeding and 0.47 from thalamic-seeding which were achieved by the true model; statistically outside the 95% confidence interval, or p < 0.05. Hence, the reported insula and thalamic-seeded NDM outperforms all simulated models’ prediction and is unlikely to be explained by chance.

**Figure 6:**
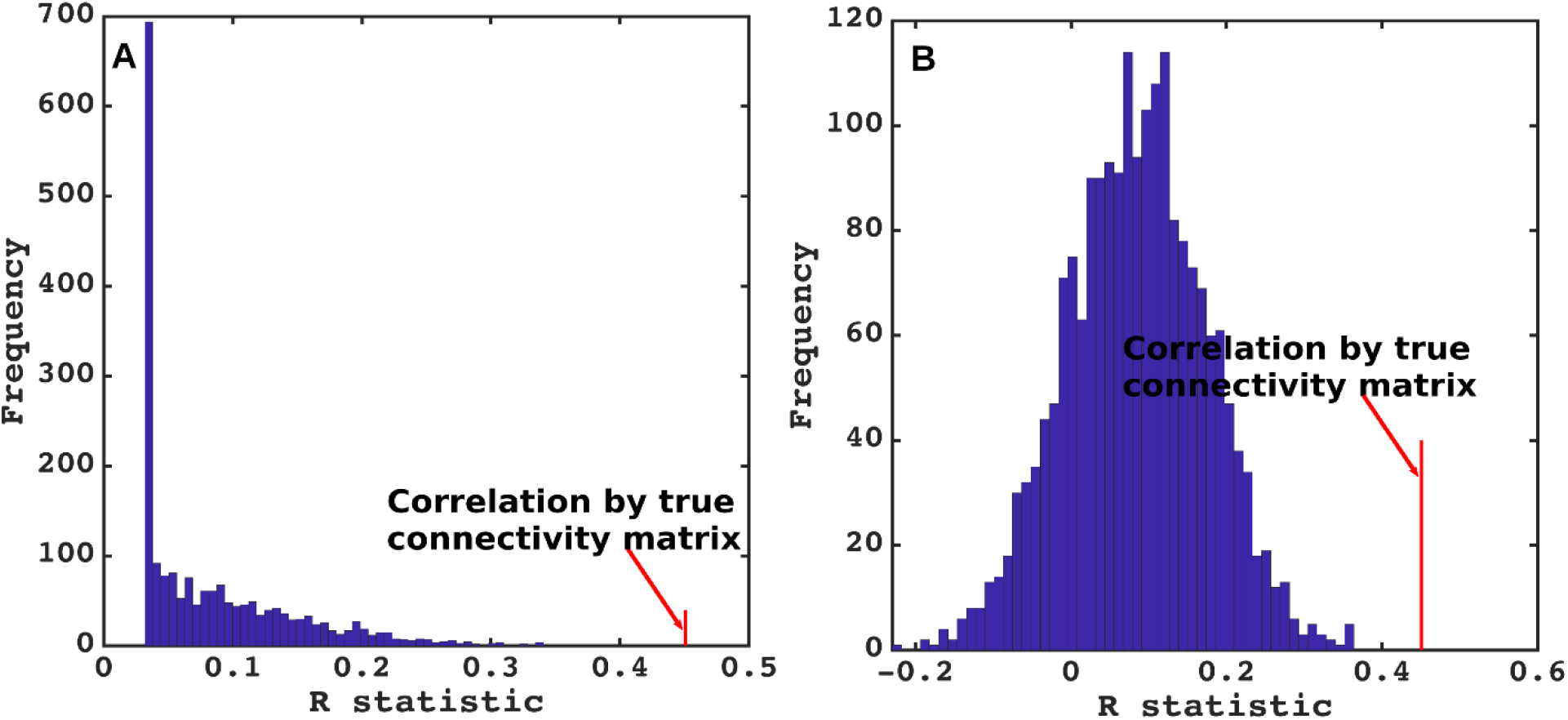
NDM evaluation against alternate models. A] Histogram of correlation strength between NDM and ALS data over 2000 shuffled networks. There is a hard limit on the left of this plot at R~0.03, which corresponds to the zero-diffusion time value of curve in Figure 1D. B] Histogram of correlation strength between NDM and 2000 shuffled ALS data over using unshuffled structural connectome. The true connectome was shuffled by symmetrically permuting its rows and columns randomly, and the NDM was evaluated for each shuffled network after bilateral insula-seeding. The best R achieved by each model was recorded and entered into the histogram. The null models are distributed well below the true model, indicating that the latter is highly unlikely to arise by chance (p < 0.05).

## Discussion

Using a quantitative network-based model of pathology spread, this study sought to explore selective vulnerability and pathological progression in the ALS brain. We tested whether, setting each region of our brain atlas as the initiation site, the subsequent network spread modeled by the NDM correctly and significantly recapitulates cross-sectional patterns of regional atrophy and *post mortem* pathology staging. We also incorporated in our model the regional expression of ALS-related genes. The results support structural network-based transmission in relation to regional atrophy, but with no significant relationship to the spatial distribution of the regional expression of ALS-related genes. Intriguingly, the critical seed regions for spread within the model were not within the primary motor cortex but in basal ganglia, thalamus and insula. NDM applied to these seed regions also recapitulated the *post mortem* histopathological staging system. Within a continuous ALS-FTD clinicopathological spectrum, these non-primary motor structures may be the sites of some of the earliest cerebral pathology.

### Stereotyped Models of anatomical spread in ALS

The focality of initial symptom onset and the non-random, typically regionally contiguous spread of symptoms in ALS was shown to be mirrored, for limb involvement at least, by spinal cord histopathology (Ravits *et al*., 2007a, b). These data were used to infer a *theoretical* model of simultaneous cerebral focal onset and spread (Ravits and La Spada, 2009), but remain unproven. The most consistent regions of cerebral pathological involvement in ALS have been the corticospinal tract and corpus callosum (Filippini *et al*., 2010; Müller *et al*., 2016), but with wider extra-motor involvement at baseline and a variable extent of both grey and white matter changes in longitudinal studies (Menke *et al*., 2014).

*Post mortem* studies defined a variably overlapping extent of TDP-43 pathology, arbitrarily divided into four “stages”, with presumed but as yet unproven sequential trans-axonal progression *in vivo* (Brettschneider *et al*., 2013). In that model initial lesions were said to develop in the agranular motor cortex, in the bulbar and spinal somatomotor neurons, and the brainstem motor nuclei (stage 1). The next affected regions were the prefrontal neocortex, the brainstem reticular formation, the pre-cerebellar nuclei, and the red nucleus (stage 2), then striatum and into the prefrontal/postcentral cortices (stage 3), finally involving anteromedial portions of the temporal lobe and the hippocampus (stage 4). This pathological staging has been supported by the same group in analysis of cross-sectional *in vivo* MRI data (Gorges *et al*., 2018).

### Role of network transmission in ALS

A plausible explanation for these patterns of progression may be trans-neuronal transmission of underlying pathology, which has been hypothesized in other neurodegenerative disorders including Alzheimer’s Disease, Frontotemporal Dementia (FTD), Parkinson’s Disease, Huntington’s Disease and Creutzfeldt–Jakob disease (Spillantini *et al*., 1998; Lee *et al*., 2001; Neumann *et al*., 2006; Hansen *et al*., 2011; Herrera *et al*., 2011; Jack and Holtzman, 2013; Jucker and Walker, 2013; Walker *et al*., 2013; Maniecka and Polymenidou, 2015; Freeze *et al*., 2020). Concepts of seeding and self-templating of aberrant, aggregate-prone proteins have extended to ALS (Polymenidou and Cleveland, 2011), with a similar hypothesis of trans-neuronal transmission of pathogenic proteins between cells (Schmidt *et al*., 2016; Subramaniam, 2019). These emerging concepts in ALS and FTD were comprehensively reviewed recently (Riku, 2020). Broader concepts of structural and functional networks in health have been invoked for defining patterns of neurodegeneration (Seeley *et al*., 2009). MRI studies in ALS have supported the concept that structural connectivity mediates the spatial and temporal evolution of ALS atrophy and leads to network disintegration (Verstraete *et al*., 2011, 2014; Schmidt *et al*., 2016; Bede *et al*., 2018).

The present study is different from previous network studies in ALS because it does not investigate the question of whether structural connectivity networks themselves are being damaged by the disease process, but whether they serve as conduits for pathology transmission on the network. (Bede *et al*., 2018) found that the subcortical areas that undergo the highest cell loss in ALS are dictated by their connectivity to cortical regions. This could be due to network degeneration as above, but could also point to protein transmission along connections, as explored in our study.

### Insula, basal ganglia and thalamus as potential seeding sites in ALS

The critical seed regions for widespread pathological spread within our model were not in the motor cortex but in basal ganglia, thalamus and insula. Furthermore, NDM applied to these seed regions recapitulated the *post mortem* TDP43-based histopathological staging scheme. These brain regions are well connected with prominent cortical areas undergoing atrophy in ALS (Bede *et al*., 2018). Basal ganglia involvement in ALS has been increasingly recognized (Bede *et al*., 2013; Riku, 2020) and the thalamus in particular has been shown to reflect the wider extent of cortical involvement in ALS (Chipika *et al*., 2020), notably in relation to the longitudinal spread of frontotemporal involvement (Tu *et al*., 2018).

A speculative interpretation of our results is that such sites may be common ‘anchors’ for what is a continuous ALS-FTD spectrum; see e.g. a recent review (Riku, 2020). The frontoinsula region appears to be one of the more selectively vulnerable and perhaps earliest sites of pathology in behavioral variant FTD, from which large von Economo neurons are prominently lost (Seeley *et al*., 2008; Kim *et al*., 2012). Similarly, the striatum is a site of early and prominent atrophy in bvFTD (Halabi *et al.*). Pathology in these non-primary motor deep gray matter structures may progress into either predominantly motor areas in ALS patients, or frontotemporal regions in FTD. Carriers of the intronic hexanucleotide expansion in *C9orf72*, the commonest inherited form of both ALS and FTD, tend to dichotomize into a phenotype with a predominance of one or other condition, even within the same pedigree (Mahoney *et al*., 2012), and the application of the methodology to a large cohort of such individuals might strengthen the hypothesis.

### Limitations

Our study was not able to accommodate many potential disease mechanisms like oxidative stress, mitochondrial damage, metabolic dysregulation, and cell-type-specific factors, e.g. the specific role of motoneurons. The NDM is a first-order, linear model of diffusive spread that assumes that the structural connectivity network remains unchanged during disease course. Although all neurodegenerative diseases lead to aberrant structural connectivity, in practice normative connectomes as used here usually do not lead to significant reduction in the model’s predictive power (Powell et al, 2017). Individual subjects’ genetic variables, medication history and age of symptom onset were not analyzed.

## ACKNOWLEDGEMENTS

Authors convey grateful thanks to Chris Mezias and Justin Torok at Weill-Cornell, and Pablo Damasceno at UCSF for help with network and gene analysis.

## Figure legends

**Figure 7: Spatial distribution of ALS atrophy and repeated seeding.** A] Measured regional ALS atrophy are depicted by glass brain visualization. Bilateral volume loss was observed in somatosensory, frontotemporal, and subcortical regions, with most atrophy occurring in precentral gyrus, inferior temporal gyrus, precuneus, putamen, and thalamus regions. Severity of disease in each region is depicted in a color bar, where color towards red show increased severity. B] Each region was seeded in turn and NDM was played out for all time points. Pearson’s R was recorded at each time point between the model and ALS atrophy vector. As the diffusion time increases, more and more of the pathogenic agent escapes the seed region and enters the rest of the network. The point of maximum correlation with measured atrophy was recorded with glass brains of measured R with spheres placed at the centroid of each brain region, and their diameter proportional to effect size. Spheres are color coded by lobe – frontal = purple, parietal = red, occipital = orange, temporal = cyan, subcortical = green, and cerebellum = magenta. C] Histogram of empirical atrophy and seed region likelihood as represented by *R_max_* is shown side-by-side. Precentral which is the highest atrophied region when taking the average of empirical atrophy from left hemisphere (LH) and right hemisphere (RH) is not the best seed, thereby suggestive of inconsequential role of higher atrophy values in determination of *R_max_*. D] NDM seeded at bilateral regions indicates that the thalamus is the one of the most plausible candidate for ALS seeding – it has the highest peak R, and the characteristic intermediate peak indicative of true pathology spread. Other regions among the top five that obtained the highest R were insula, pallidum, putamen, and caudate. R-t curves for the remaining regions are shown in blue. E] Histogram of maximum R achieved from six major regions for all individual subjects. Rmax values were attained for each of these regions from 79 individual subjects. We can see that for most of the subjects’ maximum R (Rmax > 0.4) was achieved from the frontal and subcortical regions compared to other regions.

**Figure 8: Spatial distribution of ALS atrophy, NDM Predicted atrophy and histopathological staging.** A] Measured regional ALS atrophy are depicted by glass brain visualization. Bilateral volume loss was observed in somatosensory, frontotemporal, and subcortical regions, with most atrophy occurring in precentral gyrus, inferior temporal gyrus, precuneus, putamen, and thalamic regions. Severity of disease in each region is depicted in a color bar, where color towards red show increased severity. B] Glass brains of NDM seeded at the bilateral insula at *t_max_* = 6.06 au yields progression of ALS from insula to connected, subcortical, anteromedial portions of temporal lobe and frontal areas. Bilateral volume loss is mainly observed in frontal and subcortical regions, with most atrophy occurring in later orbito-frontal, superior frontal, precentral, rostral middle-frontal, and putamen regions. Severity of disease in each region is depicted in a color bar, where color towards magenta showing increased severity. C] The ALS stage from 1-4 for each of the 43 bilateral regions. Stage 1 (maroon) starts with agranular motor cortex. The next affected regions (stage 2 in red) are the premotor cortex and parts of prefrontal neocortex. The pathology then progresses into striatum and into the prefrontal/postcentral cortices (stage 3 in orange), and finally to stage 4 show (yellow) involving anteromedial portions of the temporal lobe and the hippocampus. Stage 5 shows regions in white that are not part of the published histopathological staging system.

**Figure 9: Spatiotemporal evolution, scatter plots of atrophy and histopathological staging with NDM at different model times.** A] Evolution of insula-seeded network diffusion at model times *t* = 2,4,6 (*au*) exhibited frontal involvement initially, followed by slower diffusion into the temporal, motor and subcortical cortices, and finally showing widespread involvement of cortical regions. This temporal sequencing predicted by the model suggest that volume loss in ALS involves extra-motor regions, particularly the prefrontal and subcortical regions. B] Scatter plot of NDM from insula versus empirical ALS atrophy at model times *t* = 2,4,6 (*au*). Dots are color coded by lobe - frontal = purple; parietal = red; occipital = orange; temporal = cyan; subcortical = green; and cerebellum = magenta. A positive correlation is observed between ALS empirical atrophy and NDM predicted atrophy from bilateral insula, which increases significantly (R = 0.45, p_corr_ < 5.8×10^−4^, 0.05/86) at matured model times *t* = 4,6 (*au*). C] Scatter plot of NDM from insula versus histopathological staging at model times *t* = 2,4,6 (*au*). Dots are color coded by lobe - frontal = purple; parietal = red; occipital = orange; temporal = cyan; subcortical = green; and cerebellum = magenta. A negative correlation was observed between the NDM and ALS staging from bilateral insula, which decreases significantly (R = −0.66, p_corr_ < 5.8×10^−4^) at matured model times *t* = 4,6 (*au*). As time progressed, greater frontal, temporal and subcortical regions were involved with NDM closely resembling empirical ALS-FTD pathology.

**Figure 10: Spatial distribution of ALS atrophy, scatter plots of genes vs ALS atrophy, spatial distribution of genes.** A] Measured regional ALS atrophy are depicted by glass brain visualization. Bilateral volume loss was observed in somatosensory, frontotemporal, and subcortical regions, with most atrophy occurring in precentral gyrus, inferior temporal gyrus, precuneus, putamen, and thalamic regions. Severity of disease in each region is depicted in a color bar, where color towards red show increased severity. B] Scatter plot of empirical atrophy vs average of all ALS-related genes shows no clear association. C] Scatter plot of empirical atrophy vs average of TDP-43 associated genes (F) shows no clear association. D] Scatter plot of empirical atrophy vs TARDBP gene expression (G) also shows no clear association – this was chosen for comparison, given that a small minority of ALS cases involve mutations in TARDBP. Spheres in glass brains were placed at the centroid of each brain region, and their diameter was proportional to effect size.

**Figure 11: Lasso plots and model parameters.** A] Ten-fold cross validated MSE curves for determining regularized parameter lambda. Predictors with minimum L_1_ coefficient as a function of regularized parameter lambda with no more than one standard deviation (blue dotted line) were considered to be the most favorable. B] Cross-validated L_1_ regularized regression coefficients as a function of tuning parameter lambda for a model containing the NDM from insula, ALS-related genes, and genes implicated in trans-synaptic TDP-43 transfer as predictors. Trace plot shows that as lambda increases towards the left, lasso sets various coefficients to zero, thereby removing them from the model. C] Model parameters and p-values of significant predictors that survived with p < 0.05 (represented with “*”) and with Bonferroni corrected p (represented with “**”). The NDM and expression profiles of CCNF, UBQLN2, PFN1, and CSNK1E have non-zero coefficients at minimum model MSE, indicating that these are essential predictive variables.

**Figure 12: NDM evaluation against alternate models.** A] Histogram of correlation strength between NDM and ALS data over 2000 shuffled networks. There is a hard limit on the left of this plot at R~0.03, which corresponds to the zero-diffusion time value of curve in Figure 1D. B] Histogram of correlation strength between NDM and 2000 shuffled ALS data over using unshuffled structural connectome. The true connectome was shuffled by symmetrically permuting its rows and columns randomly, and the NDM was evaluated for each shuffled network after bilateral insula-seeding. The best R achieved by each model was recorded and entered into the histogram. The null models are distributed well below the true model, indicating that the latter is highly unlikely to arise by chance (p < 0.05).

---

**Figure SI - 1:**
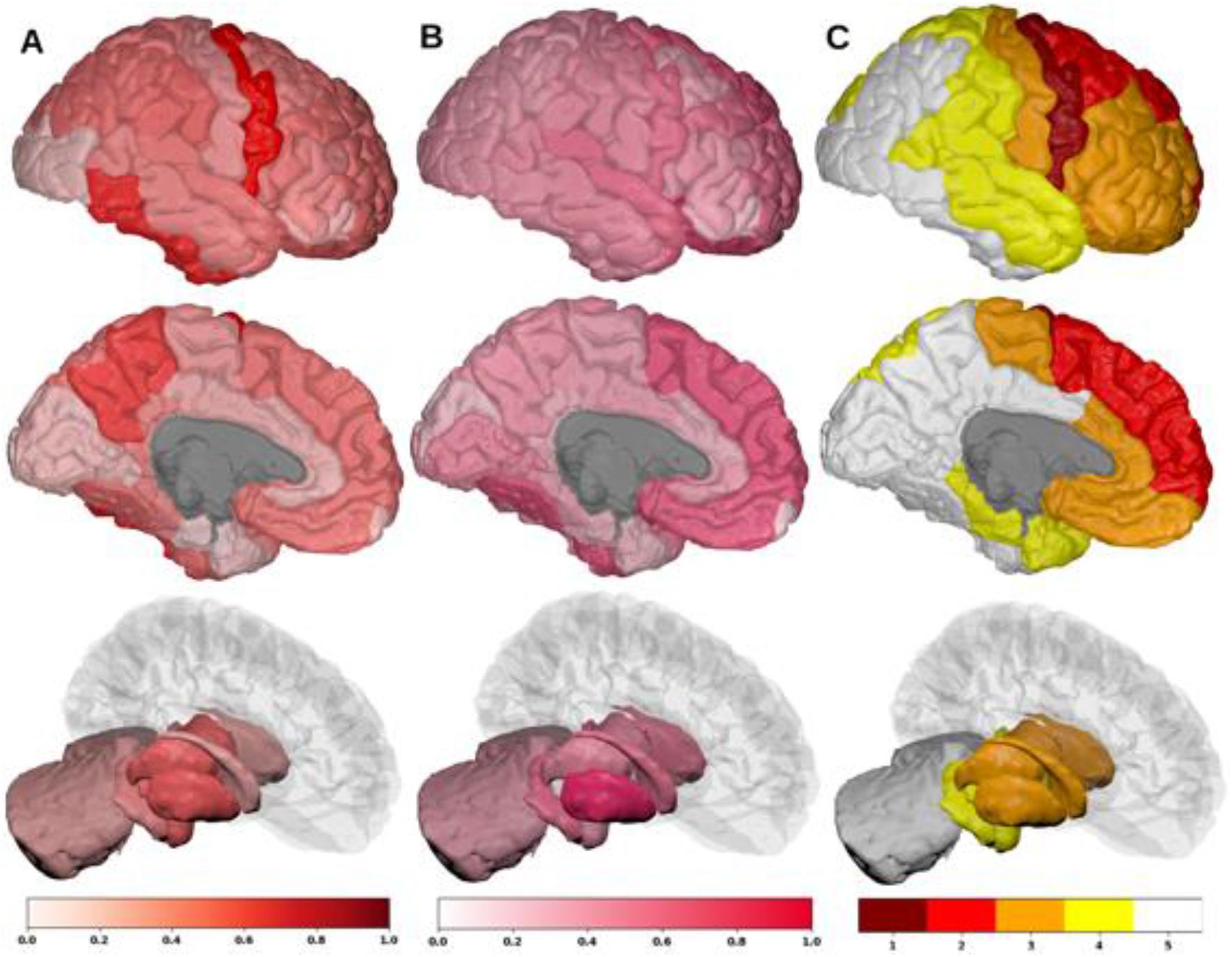
Spatial distribution of ALS atrophy, NDM Predicted atrophy and histopathological staging. A] Measured regional ALS atrophy are depicted by “glass brain” visualization. Bilateral volume loss is observed in somatosensory, frontotemporal, and subcortical regions, with most atrophy occurring in precentral gyrus, inferior temporal gyrus, precuneus, putamen, and thalamic regions. Severity of disease in each region is depicted in a color bar, where color towards red show increased severity. B] Glass brains of NDM seeded at the bilateral thalamus at *t_max_* = 8.08 au yields progression of ALS from thalamus to connected extra-motor, subcortical and frontal areas. Bilateral volume loss is mainly observed in frontal and subcortical regions, with most atrophy occurring in insula, putamen, later orbito-frontal, superior frontal, and fusiform. Severity of disease in each region is depicted in a color bar, where color towards magenta showing increased severity. C] The ALS stage from 1-4 for each of the 43 bilateral regions. We can see that Stage 1 which is indicated by maroon starts with agranular motor cortex. The next affected regions (stage 2 in red) are the premotor cortex and parts of prefrontal neocortex. The pathology then progresses into striatum and into the prefrontal/postcentral cortices (stage 3 in orange), and finally to stage 4 show in yellow involving anteromedial portions of the temporal lobe and the hippocampus. Stage 5 denotes regions in white that are not part of ALS staging schema (i.e. outside the 4 accepted stages in ALS).

**Figure SI - 2:**
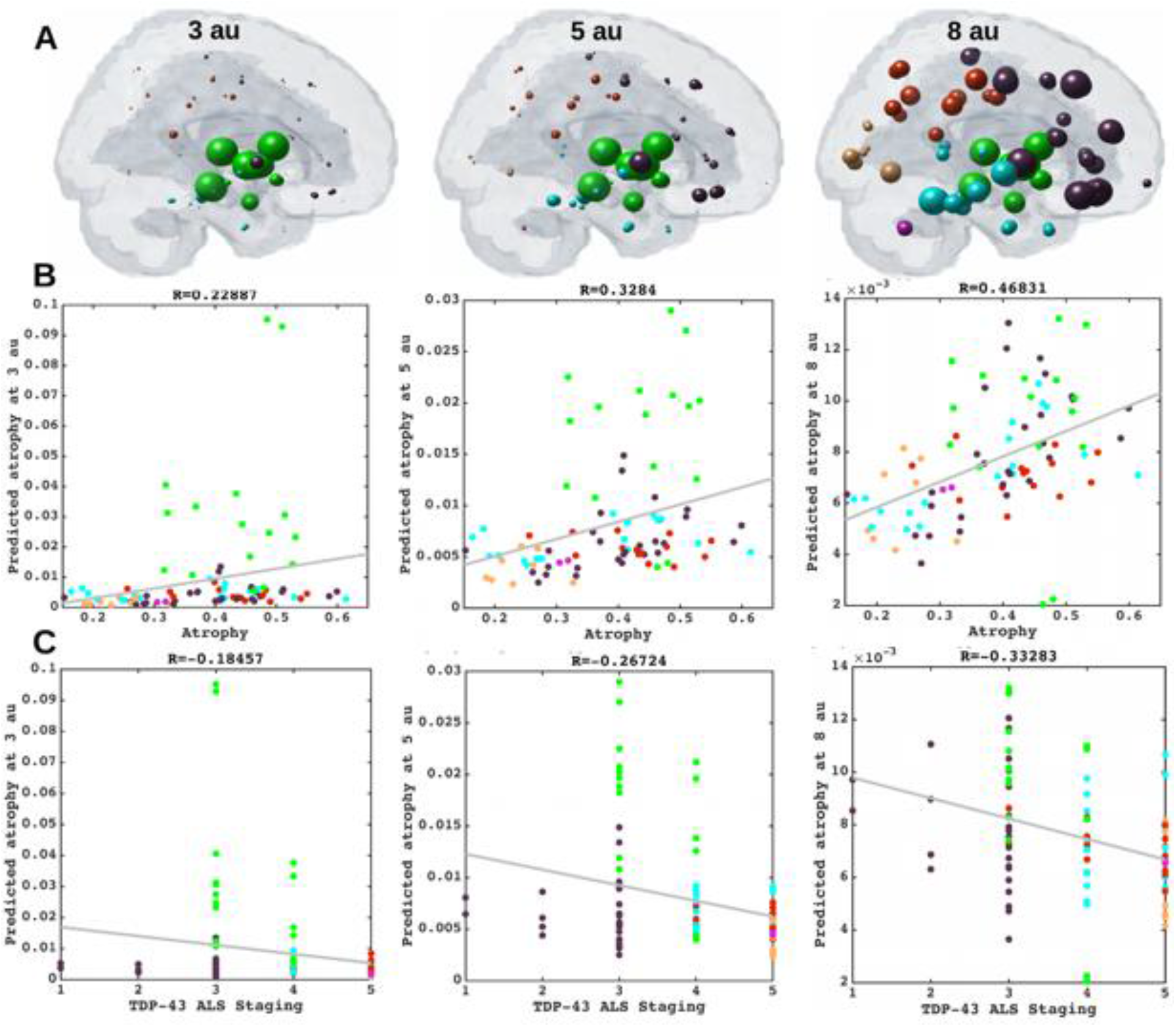
Spatiotemporal evolution, scatter plots of atrophy and histopathological staging with NDM at different model times. A] Evolution of thalamic-seeded network diffusion at model times *t* = 3,5,8 (*au*) exhibits subcortical areas as early affected regions, followed by somewhat slower diffusion into the motor and extra-motor cortices, specially prefrontal, and finally showing widespread involvement of cortical regions. This temporal sequencing predicted by the model suggest that volume loss in ALS involves extra-motor regions, particularly the prefrontal and subcortical regions. B] Scatter plot of NDM from thalamus versus empirical ALS atrophy at model times *t* = 3,5,8 (*au*). Dots are color coded by lobe - frontal = purple; parietal = red; occipital = orange; temporal = cyan; subcortical = green; and cerebellum = magenta. A positive correlation is observed between ALS empirical atrophy and NDM predicted atrophy from bilateral thalamus, which increases significantly (R = 0.47, p_corr_ < 5.8×10^−4^, 0.05/86) at matured model times *t* = 5,8 (*au*). As time progresses, more and more frontal, temporal and extra-motor regions are involved with NDM closely resembling ALS atrophy. C] Scatter plot of NDM from thalamus versus ALS staging of TDP-43 pathology at model times *t* = 3,5,8 (*au*). Dots are color coded by lobe - frontal = purple; parietal = red; occipital = orange; temporal = cyan; subcortical = green; and cerebellum = magenta. A negative correlation is observed between the NDM and ALS staging from bilateral thalamus, which decreases significantly (R = −0.33, p_corr_ < 5.8×10^−4^) at matured model times *t* = 5,8 (*au*). As time progresses, more and more frontal, temporal and subcortical regions are involved with NDM closely resembling empirical ALS atrophy.

**Figure SI - 3:**
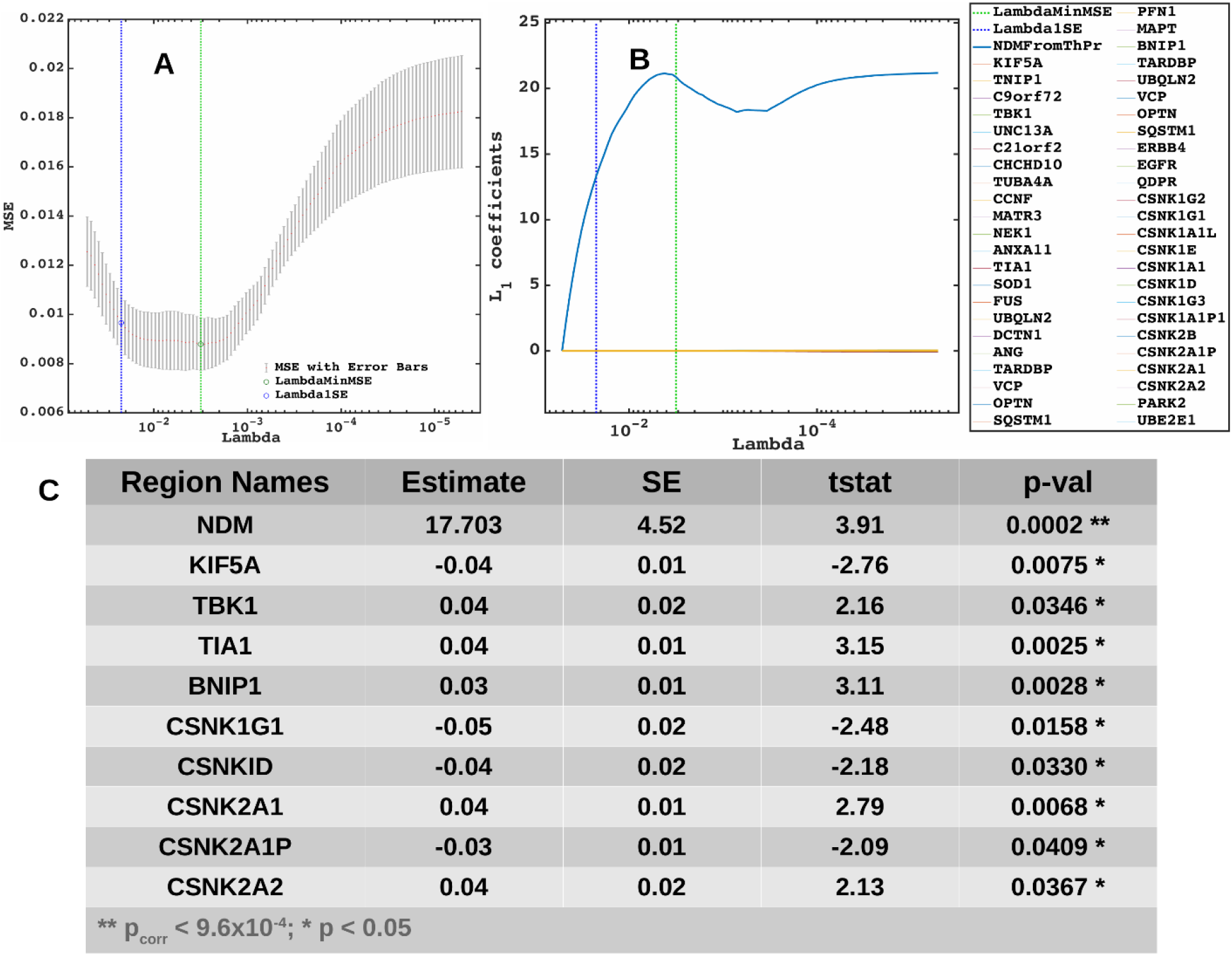
Lasso plots and model parameters. A] Ten-fold cross validated MSE curves for determining regularized parameter lambda. Predictors with minimum L_1_ coefficient as a function of regularized parameter lambda with no more than one standard deviation (blue dotted line) were considered to be the most favorable. B] Cross-validated L_1_ regularized regression coefficients as a function of tuning parameter lambda for a model containing the NDM from thalamus, ALS-related genes, and genes implicated in trans-synaptic TDP-43 transfer as predictors. Trace plot shows that’s as lambda increases towards the left, lasso sets various coefficients to zero, thereby removing them from the model. C] Model parameters and p-values of significant predictors that survived with p < 0.05 (represented with “*”) and with Bonferroni corrected p (represented with “**”). The NDM and expression profiles of KIF5A, TBK1, TIA1, BNIP1, CSNK1G1, CSNKID, CSNK2A1, CSNK2A1P, and CSNK2A2 have non-zero coefficients at minimum model MSE, indicating that these are essential predictive variables.

**Figure SI - 4:**
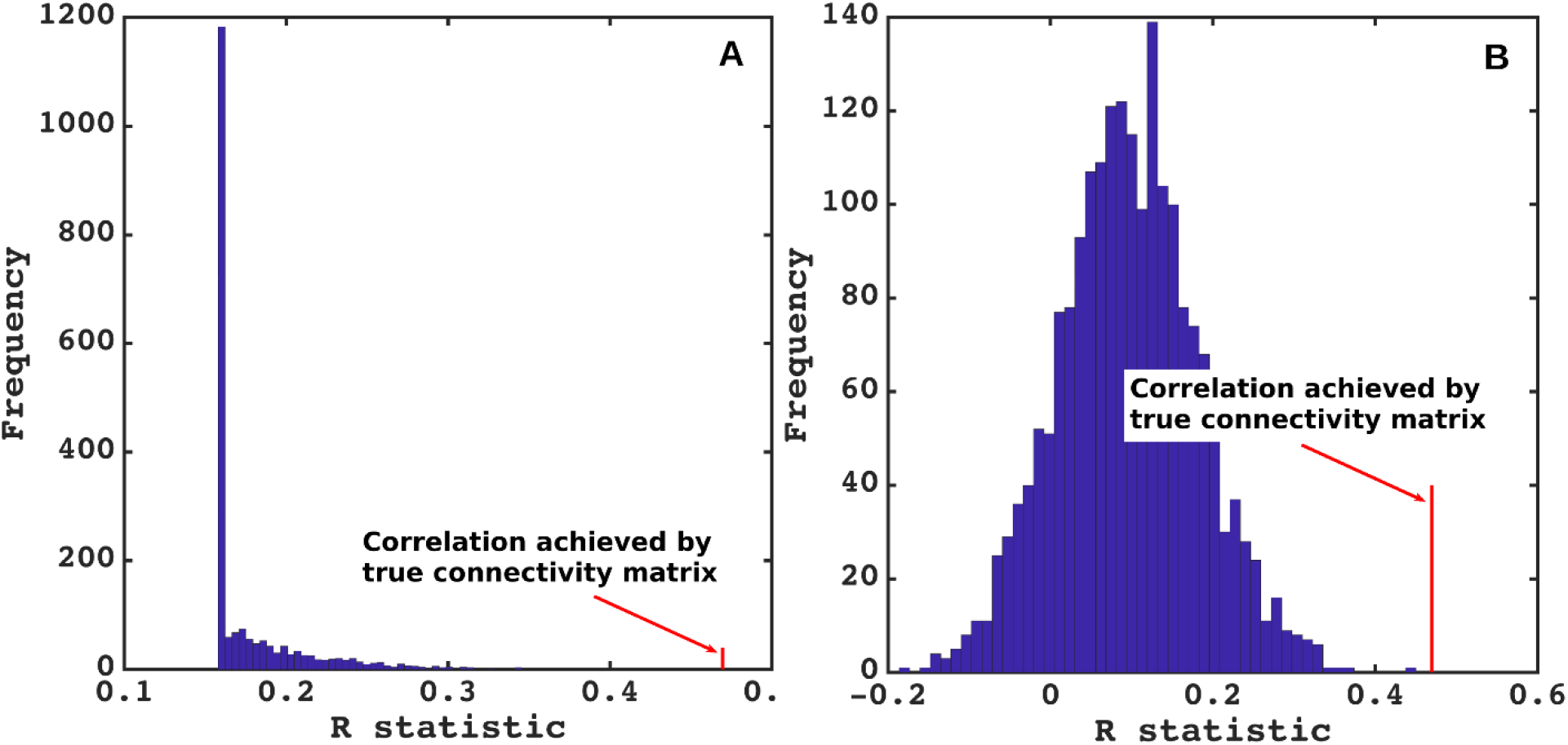
NDM evaluation against alternate models. A] Histogram of correlation strength between NDM and ALS data over 2000 shuffled networks. There is a hard limit on the left of this plot at R~0.15, which corresponds to the zero-diffusion time value of curve in Figure 1D. B] Histogram of correlation strength between NDM and 2000 shuffled ALS data over using unshuffled structural connectome. The true connectome was shuffled by symmetrically permuting its rows and columns randomly, and the NDM was evaluated for each shuffled network after bilateral thalamic-seeding. The best R achieved by each model was recorded and entered into the histogram. The null models are distributed well below the true model, indicating that the latter is highly unlikely to arise by chance (p < 0.05).

## Supplementary Information

**Figure SI - 5: Spatial distribution of ALS atrophy, NDM Predicted atrophy and histopathological staging.** A] Measured regional ALS atrophy are depicted by “glass brain” visualization. Bilateral volume loss is observed in somatosensory, frontotemporal, and subcortical regions, with most atrophy occurring in precentral gyrus, inferior temporal gyrus, precuneus, putamen, and thalamic regions. Severity of disease in each region is depicted in a color bar, where color towards red show increased severity. B] Glass brains of NDM seeded at the bilateral thalamus at t_max_ = 8.08 au yields progression of ALS from thalamus to connected extra-motor, subcortical and frontal areas. Bilateral volume loss is mainly observed in frontal and subcortical regions, with most atrophy occurring in insula, putamen, later orbito-frontal, superior frontal, and fusiform. Severity of disease in each region is depicted in a color bar, where color towards magenta showing increased severity. C] The ALS stage from 1-4 for each of the 43 bilateral regions. We can see that Stage 1 which is indicated by maroon starts with agranular motor cortex. The next affected regions (stage 2 in red) are the premotor cortex and parts of prefrontal neocortex. The pathology then progresses into striatum and into the prefrontal/postcentral cortices (stage 3 in orange), and finally to stage 4 show in yellow involving anteromedial portions of the temporal lobe and the hippocampus. Stage 5 denotes regions in white that are not part of ALS staging schema (i.e. outside the 4 accepted stages in ALS).

**Figure SI - 6: Spatiotemporal evolution, scatter plots of atrophy and histopathological staging with NDM at different model times.** A] Evolution of thalamic-seeded network diffusion at model times *t* = 3,5,8 (*au*) exhibits subcortical areas as early affected regions, followed by somewhat slower diffusion into the motor and extra-motor cortices, specially prefrontal, and finally showing widespread involvement of cortical regions. This temporal sequencing predicted by the model suggest that volume loss in ALS involves extra-motor regions, particularly the prefrontal and subcortical regions. B] Scatter plot of NDM from thalamus versus empirical ALS atrophy at model times *t* = 3,5,8 (*au*). Dots are color coded by lobe - frontal = purple; parietal = red; occipital = orange; temporal = cyan; subcortical = green; and cerebellum = magenta. A positive correlation is observed between ALS empirical atrophy and NDM predicted atrophy from bilateral thalamus, which increases significantly (R = 0.47, p_corr_ < 5.8×10^−4^, 0.05/86) at matured model times *t* = 5,8 (*au*). As time progresses, more and more frontal, temporal and extra-motor regions are involved with NDM closely resembling ALS atrophy. C] Scatter plot of NDM from thalamus versus ALS staging of TDP-43 pathology at model times *t* = 3,5,8 (*au*). Dots are color coded by lobe - frontal = purple; parietal = red; occipital = orange; temporal = cyan; subcortical = green; and cerebellum = magenta. A negative correlation is observed between the NDM and ALS staging from bilateral thalamus, which decreases significantly (R = −0.33, p_corr_ < 5.8×10^−4^) at matured model times *t* = 5,8 (*au*). As time progresses, more and more frontal, temporal and subcortical regions are involved with NDM closely resembling empirical ALS atrophy.

**Figure SI - 7: Lasso plots and model parameters.** A] Ten-fold cross validated MSE curves for determining regularized parameter lambda. Predictors with minimum L_1_ coefficient as a function of regularized parameter lambda with no more than one standard deviation (blue dotted line) were considered to be the most favorable. B] Cross-validated L_1_ regularized regression coefficients as a function of tuning parameter lambda for a model containing the NDM from thalamus, ALS-related genes, and genes implicated in trans-synaptic TDP-43 transfer as predictors. Trace plot shows that’s as lambda increases towards the left, lasso sets various coefficients to zero, thereby removing them from the model. C] Model parameters and p-values of significant predictors that survived with p < 0.05 (represented with “*”) and with Bonferroni corrected p (represented with “**”). The NDM and expression profiles of KIF5A, TBK1, TIA1, BNIP1, CSNK1G1, CSNKID, CSNK2A1, CSNK2A1P, and CSNK2A2 have non-zero coefficients at minimum model MSE, indicating that these are essential predictive variables.

**Figure SI - 8: NDM evaluation against alternate models.** A] Histogram of correlation strength between NDM and ALS data over 2000 shuffled networks. There is a hard limit on the left of this plot at R~0.15, which corresponds to the zero-diffusion time value of curve in Figure 1D. B] Histogram of correlation strength between NDM and 2000 shuffled ALS data over using unshuffled structural connectome. The true connectome was shuffled by symmetrically permuting its rows and columns randomly, and the NDM was evaluated for each shuffled network after bilateral thalamic-seeding. The best R achieved by each model was recorded and entered into the histogram. The null models are distributed well below the true model, indicating that the latter is highly unlikely to arise by chance (p < 0.05).

**Table SI - 1:** Empirical atrophy of top 20 regions (average of both left and right hemispheres)

| Region Names | Average empirical atrophy |
| --- | --- |
| Precentral | 0.5929 |
| Inferiortemporal | 0.5717 |
| Precuneus | 0.5451 |
| Putamen | 0.5099 |
| ThalamusProper | 0.4972 |
| Amygdala | 0.4918 |
| Lateralorbitofrontal | 0.4843 |
| Supramarginal | 0.4804 |
| Pallidum | 0.4790 |
| Hypothalamus | 0.4714 |
| Rostralmiddlefrontal | 0.4621 |
| Inferiorparietal | 0.4614 |
| Fusiform | 0.4599 |
| Superiorfrontal | 0.4509 |
| Superiorparietal | 0.4431 |
| Parsopercularis | 0.4425 |
| Superiortemporal | 0.4420 |
| Medialorbitofrontal | 0.4414 |
| Caudalmiddlefrontal | 0.4239 |
| Middletemporal | 0.4124 |

**Table SI - 2:**
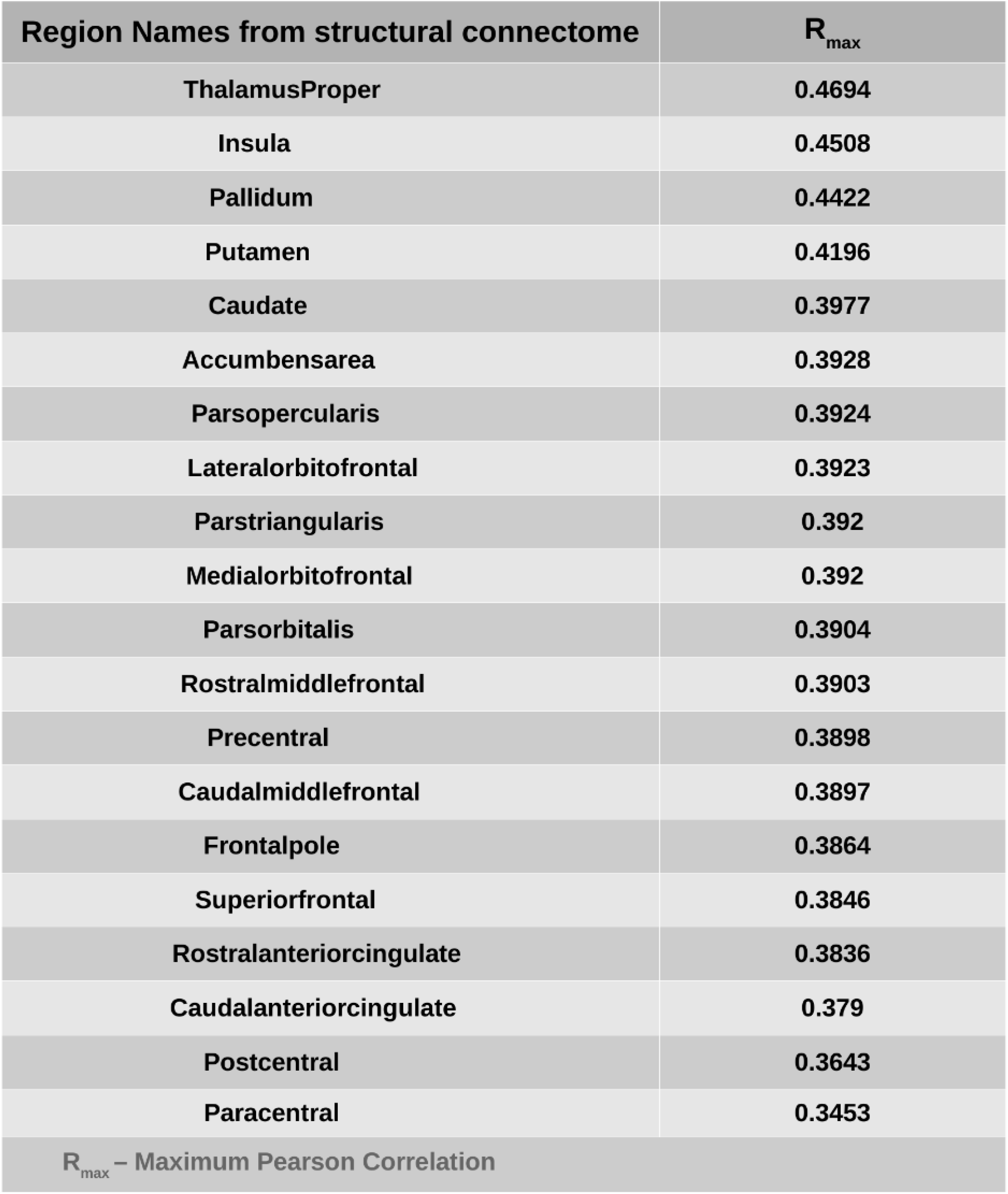
Top 20 regions with maximum correlation strength for bilaterally seeded regions

| Region Names from structural connectome | $R_{\max}$ |
| --- | --- |
| ThalamusProper | 0.4694 |
| Insula | 0.4508 |
| Pallidum | 0.4422 |
| Putamen | 0.4196 |
| Caudate | 0.3977 |
| Accumbensarea | 0.3928 |
| Parsopercularis | 0.3924 |
| Lateralorbitofrontal | 0.3923 |
| Parstriangularis | 0.392 |
| Medialorbitofrontal | 0.392 |
| Parsorbitalis | 0.3904 |
| Rostralmiddlefrontal | 0.3903 |
| Precentral | 0.3898 |
| Caudalmiddlefrontal | 0.3897 |
| Frontalpole | 0.3864 |
| Superiorfrontal | 0.3846 |
| Rostralanteriorcingulate | 0.3836 |
| Caudalanteriorcingulate | 0.379 |
| Postcentral | 0.3643 |
| Paracentral | 0.3453 |
| $R_{\max}$ – Maximum Pearson Correlation | |

**Table SI - 3:** Correlations of ALS-related genes, and their PCA vs empirical atrophy (to the left) and correlations of TDP-43 specific genes and their PCA vs empirical atrophy (to the right).

| All implicated genes |  |  | TDP-43 genes |  |  |
| --- | --- | --- | --- | --- | --- |
| Gene Names | Pearson R | p-val | Gene Names | Pearson R | p-val |
| KIF5A | -0.1656186196 | 0.1275200472 | TARDBP | 0.0587166421 | 0.5912596775 |
| TNIP1 | -0.047536623 | 0.6638298672 | UBQLN2 | -0.1408213232 | 0.1959180963 |
| C9orf72 | -0.2210541014 | 0.040202527 | VCP | -1.99E-01 | 6.65E-02 |
| TBK1 | -0.0512079554 | 0.6396119229 | OPTN | 0.2324602021 | 0.0312558543 |
| UNC13A | 0.0658038117 | 0.5471989112 | SQSTM1 | 0.046762839 | 0.6689795153 |
| C21orf2 | 0.1303410162 | 0.2316356969 | ERBB4 | 0.1651074327 | 0.1287159075 |
| CHCHD10 | 0.298541968 | 0.0052379237 | EGFR | -0.0495807645 | 0.6503011148 |
| TUBA4A | 0.1365186655 | 0.2100696947 | QDPR | 0.1376219042 | 0.206373713 |
| CCNF | 0.1614933579 | 0.1374160643 | CSNK1G2 | -0.2607516048 | 0.0153154119 |
| MATR3 | -0.2673955129 | 0.0128125354 | CSNK1G1 | -0.1431117999 | 0.1886707366 |
| NEK1 | -0.0228555052 | 0.834542713 | CSNK1A1L | 0.0437732909 | 0.6890175914 |
| ANXA11 | -0.1058693483 | 0.3319691099 | CSNK1E | -0.2381853029 | 0.0272160232 |
| TIA1 | -0.3156106649 | 0.0030756285 | CSNK1A1 | -0.059771397 | 0.5846018753 |
| SOD1 | 0.2477350679 | 0.0214620101 | CSNK1D | -0.032865878 | 0.7638668473 |
| FUS | -0.0717255375 | 0.5116516682 | CSNK1G3 | -0.1775994452 | 0.1018558544 |
| UBQLN2 | -0.1408213232 | 0.1959180963 | CSNK1A1P1 | 0.0813687883 | 0.4564062443 |
| DCTN1 | 0.128764788 | 0.2373770587 | CSNK2B | 0.2155003034 | 0.0462938926 |
| ANG | -0.0558291315 | 0.6096588566 | CSNK2A1P | 0.0962652913 | 0.3779312784 |
| TARDBP | 0.0587166421 | 0.5912596775 | CSNK2A1 | -0.0830659204 | 0.4470364806 |
| VCP | -0.1988299645 | 0.0664668969 | CSNK2A2 | 0.201434283 | 0.0629134834 |
| OPTN | 0.2324602021 | 0.0312558543 | PARK2 | -0.0576630116 | 0.5979443426 |
| SQSTM1 | 0.046762839 | 0.6689795153 | UBE2E1 | 0.1597739631 | 0.1417082943 |
| PFN1 | 0.383757023 | 0.0002648659 | UBE2E2 | -0.0478000201 | 0.6620804566 |
| MAPT | 0.07828104 | 0.4737282635 | UBE2E3 | 0.184386267 | 0.0892228173 |
| BNIP1 | 0.1176351583 | 0.2807256022 | USP14 | -0.2344893188 | 0.0297697703 |
| PCA1 | 0.1797 | 0.0979 | USP8 | 0.0853785204 | 0.4344434894 |
| PCA2 | 0.0515 | 0.6375 | PCA1 | 0.0625 | 0.5676 |
|  |  |  | PCA2 | 0.1850 | 0.0881 |

